# Extracellular Exosomal RNAs are Glyco-Modified

**DOI:** 10.1101/2024.12.17.628987

**Authors:** Sunny Sharma, Xinfu Jiao, Jun Yang, Megerditch Kiledjian

## Abstract

Epitranscriptomic modifications play pivotal roles in regulating RNA function, encompassing base alterations and the addition of both canonical m^7^G and noncanonical nucleotide metabolite caps. Recently, the spectrum of modifications has extended to include glyco modification at the 5’ cap or within the RNA. Despite this expansion, the functional implications of glyco modification on RNA remain elusive. Our study reveals that mammalian cells labeled with N-azidoacetylgalactosamine-tetraacylated (Ac4GalNAz) produce small noncoding and nonpolyadenylated glyco-modified RNA (glycoRNA), primarily localized within exosome vesicles. The resistance of glycoRNA-containing exosomes to RNase treatment suggests that Ac4GalNAz-derived glycoRNA constitutes intraluminal cargo, distinct from recently reported cell surface glycoRNAs. Furthermore, we demonstrate that exosome cargo can be transferred to naïve cells, underscoring exosome-mediated intercellular communication of glycoRNAs. Inhibition of exosome biogenesis leads to the accumulation of intracellular glycoRNA while blocking glycan transfer to proteins concomitantly reduced the targeting of glycoRNA within exosomes. These findings highlight a correlation between protein and RNA glycosylation, suggesting that the accumulation of glycoRNAs within exosomes is a regulated process. Our results support a functional role for glyco-modification in mediating RNA targeting into exosomes, offering new insights for enhancing recent advances in exosome-directed diagnostics and therapeutic applications.

## Introduction

Chemical modifications play a pivotal role in the intricate management of biomolecular sorting within cells, ensuring the precise localization of diverse molecules^1^. Extensive research has characterized modifications such as phosphorylation, acetylation, and glycosylation, providing a comprehensive understanding of their roles in guiding proteins to specific cellular compartments^1^. In stark contrast, the sophisticated language of chemical modifications governing the sorting of nucleic acids remains largely unexplored. RNA modifications have long been known to influence the fate and function of coding and noncoding RNAs. The most well-characterized modification is the 7-methylguanosine (m^7^G) cap^2^ added to the 5′ end of virtually all mRNAs, which protects the mRNA and enables ribosomal translation. Another common modification is N6-methyladenosine (m^6^A)^3^, which affects ∼30% of mRNAs and impacts mRNA stability, localization, and translation^4^. More recently, a diverse family of non-canonical caps was discovered decorating the 5′ ends of RNAs, including nucleotide metabolites like NAD, FAD, dpCoA, UDP-glucose, and UDP-N-acetylglucosamine (UDP-GlcNAc)^5^. The functions of most of these novel non-canonical caps remain unclear, except for NAD and FAD caps, which facilitate rapid RNA decay in eukaryotes^6,7^ and are employed by T4 bacteriophages in the RNAylation^8^ of bacterial translational machinery. Notably, the 5’-FAD capping^9^ of the Hepatitis C virus was demonstrated to shield the viral RNA from RIG-I-mediated innate immune recognition.

Recently the identification of glycoRNAs—small RNAs adorned with sugar molecules— on the surface of cells ^10^ has opened up new avenues for understanding RNA functionality beyond the intracellular environment. These glycoRNAs, with their distinct biochemical properties, have been proposed to play a crucial role in cellular processes, including immune response and gene expression regulation^11,12^. Our studies demonstrate the presence of glycoRNAs within extracellular exosome vesicles, suggesting a sophisticated sorting mechanism that leverages glycosylation as a determinant for exosomal packaging. This novel insight into exosomal cargo selection not only aligns with the emerging recognition of extracellular RNAs as potential biomarkers for disease but also underscores the intricate regulatory networks orchestrating RNA transport and function. We propose that the glycosylation of RNA serves as a selective signal for their inclusion into exosomes, thereby influencing the repertoire of messages conveyed between cells and potentially modulating recipient cell behavior. These findings contribute to the growing body of evidence that extracellular RNAs, particularly those associated with the plasma membrane^11^ are integral to the complex intercellular communication landscape and may hold the key to unlocking new diagnostic and therapeutic strategies.

## Results

### Identification of a distinct class of human glyco-modified small RNAs via metabolic labeling with N-Azidoacetylgalactosamine-Tetraacylated (Ac4GalNAz)

Among the recently discovered noncanonical RNA cap structures^5^, one of the most unique modifications emerges from a structurally diverse array of nucleotide sugars. Remarkably, eukaryotic cellular RNA molecules have been identified to contain 5′ uridine diphosphate N-acetyl glucosamine (UDP-GlcNAc) and UDP-glucose (UDP-Glc) moieties. Quantification via mass spectrometry has revealed a strikingly higher abundance of UDP-GlcNAc compared to the extensively studied NAD-capped RNAs^5^. To elucidate the functional role of these novel 5’-end noncanonical modifications, the identification of the decapping enzyme stands as a crucial milestone, as recently demonstrated for NAD-capped RNAs^7,13^. Considering the remarkable diversity of the substrates targeted by the DXO family of proteins^14^, we determined whether DXO and Rai1 proteins possess the capacity to hydrolyze a GlcNAc cap. In vitro, transcribed UDP-GlcNAc capped RNAs (Extended Data Figure 1a) were efficiently decapped by His-tagged recombinant Dxo or Rai1 from *E. coli*. (Extended Data Figure 1b). Consistent with their respective activities^15^, Dxo degrades the RNA following removal of the GlcNAc cap while Rai1 does not have exonuclease activity and leaves the RNA intact following UDP-GlcNAc removal (Extended Data Fig 1b). These findings demonstrate the Dxo family of proteins can hydrolyze a GlcNAc cap.

To determine whether the UDP-GlcNAc caps are detected on endogenous RNA, we leveraged and implemented the N-azidoacetylgalactosamine-tetraacylated (Ac4GalNAz) metabolic labeling approach^16^ (Figure 1a). This approach takes advantage of metabolic cross-talk between the N-acetylgalactosamine salvage and O-GlcNAcylation pathways for the tagging and identification of O-GlcNAcylated proteins^16^ and an approach analogous to a recent report for the identification of internal N-glycan modifications on small RNAs with N-azidoacetylmannosamine-tetraacylated (Ac4ManNAz)^10^ (Figure 1a). Three different glyco derivatives, Ac4GalNAz, Ac4ManNAz, and N-azidoacetylglucosamine-tetraacylated (Ac4GlcNAz) were used as potential labeling agents in HeLa cells, human embryonic kidney (HEK) 293 and breast cancer MCF7 cell lines (Figure 1a), for the identification of 5′ UDP-GlcNAc caps on cellular RNA.

**Figure 1.**
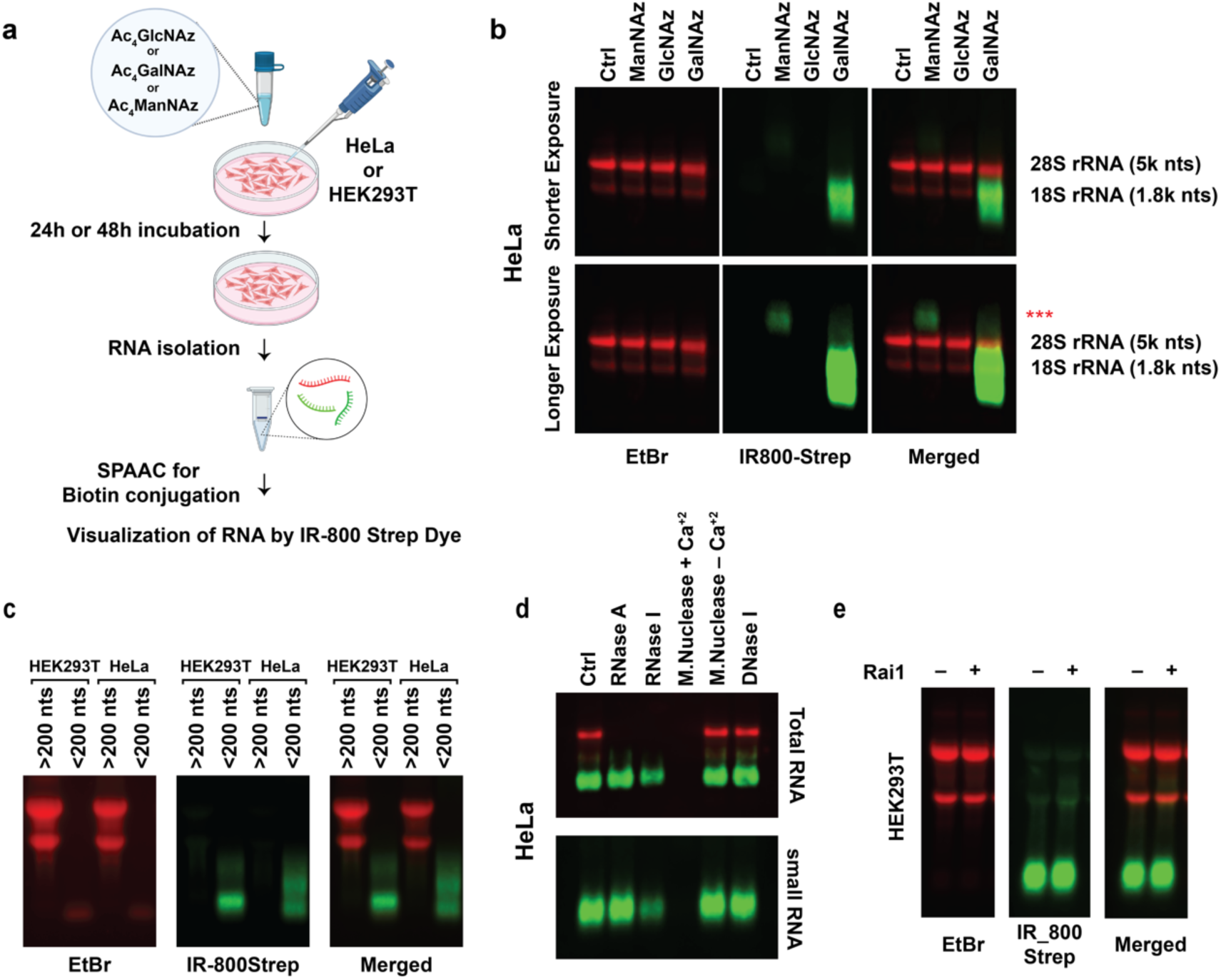
Cellular RNAs undergo sugar modifications derived from N-acetyl-galactosamine. **(a)** Experimental design schematic illustrating the characterization of labeled RNA species using biorthogonal and cell-permeable Ac_4_GlcNaz, Ac_4_GalNaz, and Ac_4_ManNaz. **(b)** Total RNA from HeLa, isolated after 48 hours of incubation with media containing 100µM of Ac_4_GlcNAz, Ac_4_GalNAz, or Ac_4_ManNAz following SPAAC reaction to conjugate biotin moieties. The resulting biotin-conjugated RNAs were resolved on an agarose gel and transferred onto a Nitrocellulose membrane. Visualization was achieved through near-infrared (IR) IRDye® 800CW streptavidin-based detection for glyco-modified RNAs and with ethidium bromide for the total RNA using Odyssey Fc (Li-Cor Biosciences). Red asterisks indicate RNAs derived from Ac_4_ManNAz, which are visible only upon extended exposure. **(c)** Ac_4_GalNaz-derived labeled total RNAs from both HeLa and HEK293T cells were fractionated into smaller (<200nts) and larger (>200nts) RNAs, followed by SPAAC reaction for biotin conjugation. The biotin-conjugated glycoRNAs were then visualized as in b. **(d)** Gel analysis depicting the impact of the indicated nucleases on the RNA. **(e)** To probe the presence of a 5’ end cap associated with the detected azide, endogenous glycoRNAs was treated with SpRai1 to remove the GlcNAc cap. Following incubation with 100µM of Ac4GalNAz A, 25µg of total RNA isolated from HEK23T cells underwent treatment with 100 nM of SpRai1. The resultant reaction product was subjected to SPAAC and visualized as in b. The lack of significant reduction in the intensity of the IR-800 signal suggests the glyco-modification likely resides internally rather than at the 5’ end.

To visualize the azide-labeled RNA, we employed strain-promoted azide-alkyne click chemistry (SPAAC) ^17,18^. Interestingly, a distinct temporal and selective Ac4GalNAz-dependent pattern in the biotinylated RNA species was evident (Figure 1b, and Extended Data Figure 2a, and 2b). Moreover, the migration of the glycoRNA on agarose gel electrophoresis indicated the modification occurred exclusively within smaller RNAs. Although no detectable biotinylated species were discerned when employing Ac4ManNAz or Ac4GlcNAz for labeling, a faint larger migrating species was detected with the Ac4ManNAz upon overexposure (Figure 1b). The presence of slower migrating glycoRNAs detected with Ac4ManNAz was consistent with a recent report^10^. Considering the prominent signal with the Ac4GalNAz labeled RNA, we focused the subsequent analysis on Ac4GalNAz glycosylated RNAs.

To further confirm the size distribution of the faster migrating GalNAz labeled RNA, the size was validated through a size-dependent RNA precipitation and silica column-based fractionation method, segregating transcripts into “large” (>200 nt) and “small” (<200 nt) categories. Consistently, glycoRNAs extensively co-fractionated with the small RNA population (Figure 1c) and were primarily devoid of polyadenylated RNA (Extended Data Figure 2c). Interestingly, the detected glycoRNAs were not sensitive to treatment with the sequence restricted nuclease, RNase A, yet sensitive to the more sequence independent RNase I nuclease where RNA degradation was evident (Figure 1d). Moreover, the signal was eliminated upon exposure to the more pleotropic nuclease, micrococcal nuclease prior to resolution on the gel with the expected Ca^2+^ dependence (Figure 2d and Extended Data Figure 2d). Consistent with the signal being derived from RNA, it was resistant to DNase treatment (Figure 1d). These findings suggest that cells labeled with Ac4depGalNAz predominantly incorporate the azide label into relatively small structured cellular RNA species.

**Figure 2.**
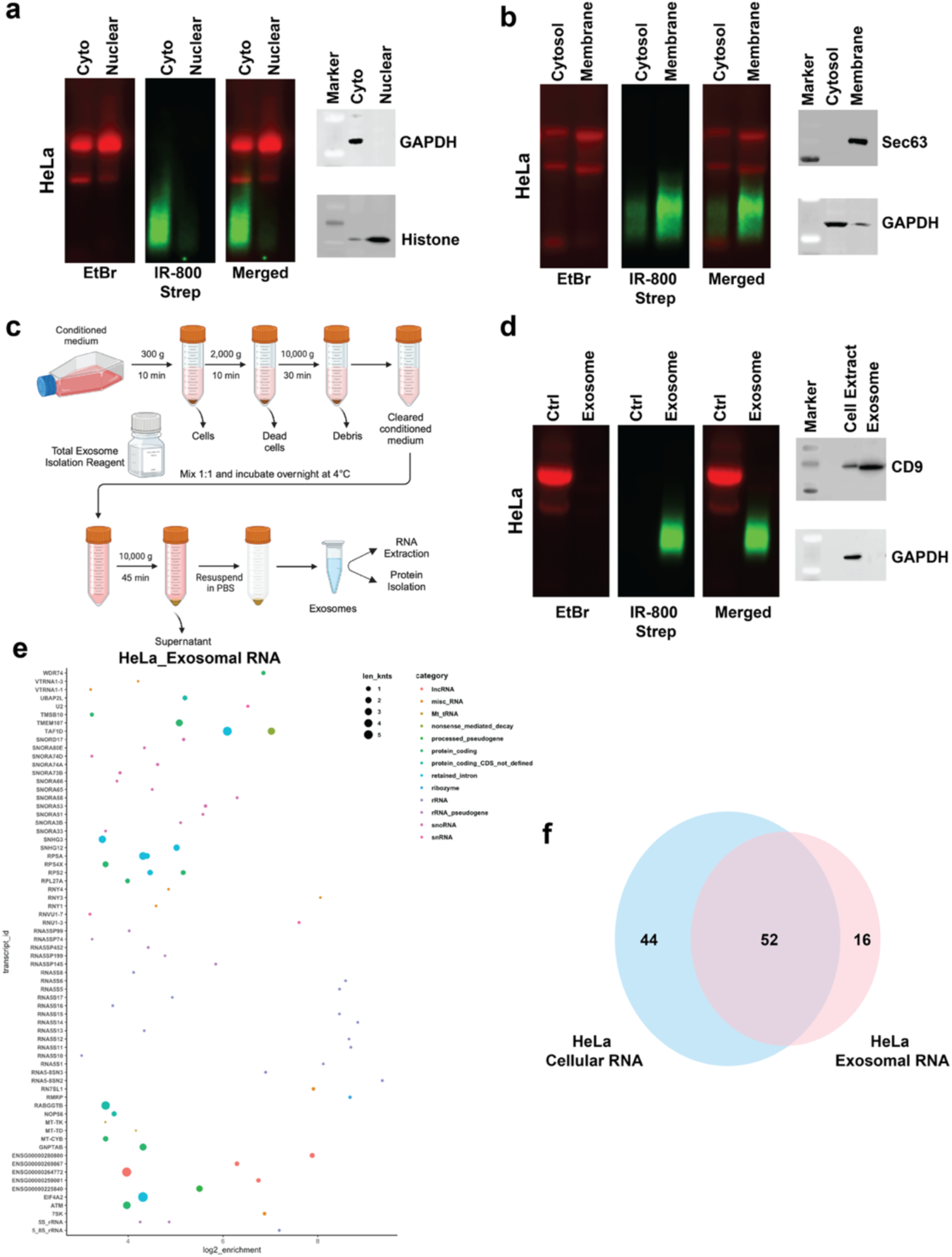
Analysis of Glyco-Modification in Exosomal RNAs. **(a)** Subcellular localization of glyco-modified RNA was evaluated in Ac_4_GalNaz-labeled HeLa cells. Nuclear and cytoplasmic fractions were resolved, RNA isolated and biotin-conjugated RNAs were visualized as in Figure 1. Western Blotting with GAPDH and histone H3 served as controls for fractionation. **(b)** Localization of glyco-modified RNAs in the soluble cytosolic compartment or membranous organelles were assessed using the ProteoExtract® Native Membrane Protein Extraction Kit (EMD Millipore). Membranous fractions (plasma membrane, ER, and Golgi) and cytosol were seperated and total RNA from each fraction were subjected to SPAAC for biotin conjugation. The biotin-conjugated glycoRNAs were analyzed as described in (a). Western Blotting with GAPDH and Sec63 served as controls for fractionation. **(c)** Schematic illustration of the exosomal RNA isolation method employed. **(d)** Exosomal RNAs derived from HeLa cells were isolated and the resulting biotin-conjugated RNAs were analyzed after SPAAC as in (a). Western Blotting with GAPDH and CD9 served as controls for exosome purification. **(e)** Scatter plot analysis depicting Ac_4_GalNaz-derived labeled exosomal RNAs from HeLa cells. **(f)** Venn diagram illustrating the overlap between cellular RNA and exosomal RNA gene sets (logFC>4), demonstrating shared and distinct transcripts.

To determine whether the azide detected on the endogenous RNA was constituted by a 5′ end cap, the RNA was treated with SpRai1 to remove the GlcNAc cap. Surprisingly, a decrease in the intensity of the IR-800 signal was not evident implying that the glyco-modification is not on the 5′ end but likely an internal modification (Figure 1e). Collectively, our findings indicate that the prevailing glycoRNA detected in HEK293T, and HeLa cells predominantly represent an internal GalNAc modification rather than a UDP-GlcNAc cap.

### Transcriptome-wide mapping of Glyco-modified RNAs identified a diverse group of small noncoding RNAs

To identify glyco-modified transcripts, small RNA fractions were first isolated from Ac4-GalNAz-labeled HeLa and HEK293T cells (Extended Data Figure 3a). Following SPAAC and affinity purification with streptavidin beads, the captured biotinylated RNAs (Extended Data Figure 3b), were next subjected to Oxford nanopore sequencing. Given that Oxford nanopore sequencing primarily targets poly(A)-containing RNAs, an additional polyadenylation step was introduced using *E. coli* poly(A) polymerase. This step was necessary since the Ac4GalNAz-labeled RNAs did not yield detectable poly(A) RNAs (Extended Data Figure 2c). The sequencing outcomes revealed a diverse collection of small noncoding RNAs, including various isoforms of snoRNAs, snRNAs, vault RNAs, and Y RNAs in both HeLa and HEK293T cells (Extended Data Figure 3c and 3d). A parallel experiment without Ac4GalNAz-labeled RNA served as a negative control, ensuring the specificity of the approach. Enrichment results were further validated by Northern blot analysis of two representative RNAs, RNY3 and 5S rRNA, using ^32^P labeled transcript-specific probes (Extended Data Figure 3e). Analysis of the identified RNAs revealed a remarkable congruence with previously identified exosomal RNAs^19,20^, including Y RNAs, vault RNAs, and rRNAs suggesting a close association between glycoRNAs and membrane organelle.

### Exosomal RNAs are modified with glyco-derived modifications

To begin unraveling the role of the glyco modifications of RNA in cellular physiology, we evaluated their intracellular localization. Two distinct biochemical approaches were utilized to determine the distribution of glycoRNAs within cellular compartments. The first involved fractionation of nuclei away from total cytosol while the second further refined the cytoplasmic fraction into the soluble cytosolic compartment and membranous organelles. IR-800 signal was almost exclusively detected in RNAs derived from Ac4GalNAz-labeled total cytoplasmic fraction but not nuclear fraction (Figure 2a). Moreover, the cytoplasmic glycoRNAs were predominantly detected within the membrane fraction relative to the soluble fraction (Figure 2b) suggesting the glycoRNAs are primarily associated with membranes. However, although Ac4MalNAz-labeled RNAs were recently shown to localize to the extracellular surface of the HeLa cell membrane^10^, the glycoRNAs described here are likely distinct from the previous report. First, micrococcal nuclease treatment of intact cells did not appreciably diminish glycoRNA detection (Extended Data Figure 4a) indicating the Ac4GalNAz-derived RNAs are not localized on the cell surface. Second, distinct categories of small RNAs (sRNAs) were recently delineated within the endoplasmic reticulum (ER)^21^ suggesting internal membranes may house the glycoRNAs. Third, the correspondence of the identified glycoRNAs (Extended Data Figures 3c and 3d) to exosome RNAs strongly support exosomal association^19,20^ (see below).

To directly determine whether glycoRNAs localized within extracellular membrane vesicles, the schematic outlined in Figure 2c was employed for exosomal RNA isolation. Fractionation of the exosome was validated utilizing Western blot analysis of GAPDH and exosome marker CD9^22^ (Figure 2d). RNA extracted from Ac4GalNAz-labeled HeLa cell exosomes were subjected to SPAAC analysis, and the resulting biotin-conjugated RNAs were evaluated. SPAAC analysis demonstrated that the Ac4GalNAz-derived glycoRNA were almost exclusively contained within HeLa cell exosomes (Figure 2d). Importantly, the glycoRNAs were not detected on the extracellular surface. Subjecting the Ac4GalNAz-labeled exosomes to micrococcal nuclease did not affect the labeled glycoRNA content (Extended Data Figure 4b) indicating that glycoRNAs are encapsulated within the exosomal lumen, rather than being surface-associated.

The composition of exosomal glycoRNAs was next addressed. Glyco-modified exosome RNAs were affinity purified using streptavidin beads, and the eluted biotinylated RNAs were subjected to Oxford nanopore sequencing (Figure 2e). A striking correlation in the RNA species was evident between the glycoRNAs from cellular and exosomal sources (Figure 2f and Extended Data Figure 4c). This finding underscores the consistency and similarity in the glyco-modification profiles between cellular and exosomal RNA populations.

To assess the relative abundance of glyco-modified RNAs in both exosomes and cells, we compared the labeling of RNA used for the SPAAC. Given that exosomes predominantly contained small RNAs (sRNAs), our focus narrowed to the small RNA fraction within cellular RNAs. Intriguingly, HeLa exosomes harbored a disproportionate percentage of labeled glycoRNAs compared to cellular sRNAs (Extended Data Figure 4d) indicting a potential functional significance for the glycosylation to target RNA to the exosome.

### ESCRT-dependent and independent pathways govern the release of glyco-modified RNA-enriched exosomes

The formation of exosomes involves an active contribution of the endosomal-sorting complex required for transport (ESCRT) machinery and a key component of this activity is the protein Hepatocyte Growth Factor-Regulated Tyrosine Kinase Substrate (HGS) also known as Vps27^23,24^ (herein referred to as HGS). A notable mechanistic dichotomy driving ESCRT-dependent and ESCRT-independent pathways exists^25^ (Figure 3a). Manumycin-A (MA)^22^, a natural microbial metabolite is a potent inhibitor of exosome biogenesis and secretion which is ESCRT-dependent with no discernible impact on cell growth. Similarly, ceramide, whose biosynthesis is regulated by neutral sphingomyelinase 2 (nSMase2) is critical for ESCRT-independent secretion of exosomes^26^, and inhibition of nSMase2 by GW4869^26^ leads to a block of exosome release.

**Figure 3.**
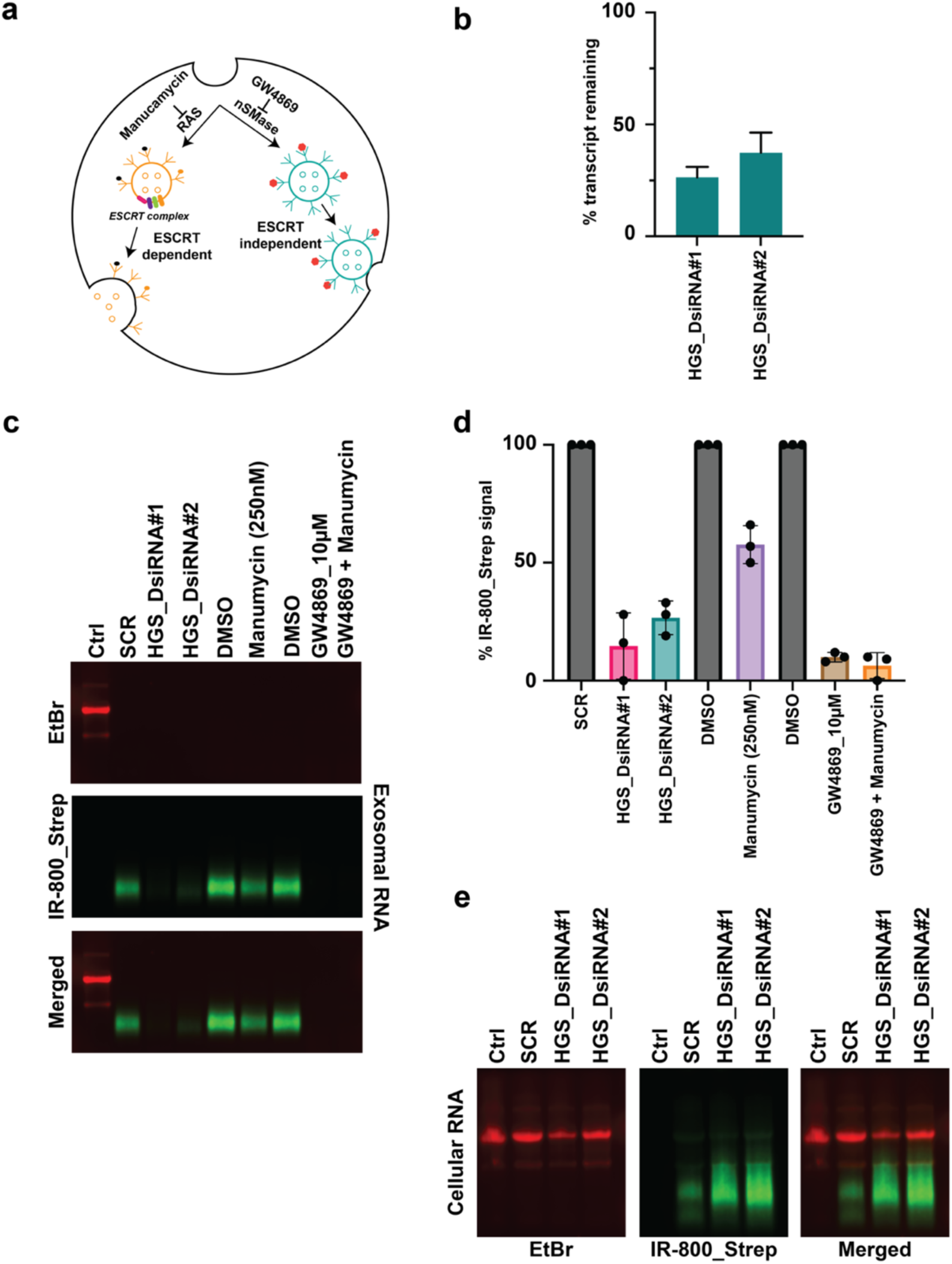
Disruption of Exosome Biogenesis/Release and Impact on Glyco-Modified RNA Dynamics. **(a)** Schematic representation of endosomes, illustrating both ESCRT-dependent and independent pathways involved in exosomal release. **(b)** HGS (Vps27) knockdown using distinct DsiRNAs. HeLa cells were transfected with two different DsiRNA targeting HGS (Vps27), followed by a 72-hour incubation. The efficiency of gene knockdown was assessed through qRT-PCR analysis of total RNA extracted from these cells. **(c)** Examination of exosomal RNAs from HGS-depleted or cells treated with Manumycin, GW4869, or a combination of Manumycin and GW4869. After 24-hour DsiRNA treatment or inhibitors, cells were exposed to 100µM Ac4GalNaz, and RNA was isolated after 48 hours of incubation with media containing 100µM Ac_4_GlcNAz. Subsequent SPAAC-mediated conjugation enabled the separation of resulting biotin-conjugated RNAs on an agarose gel and visualized as in Figure 1. **(d)** Representation of the average signal from glyco-modified RNA (n = 3) independent experiments, with error bars denoting ±SEM. **(e)** Analysis of cellular RNAs from the same set of HGS knockdown experiments (in panel c). After SPAAC-mediated conjugation, resulting biotin-conjugated RNAs were separated on an agarose gel and visualized as in Figure 1.

To substantiate the presence of glycoRNAs in exosomes and examine the influence of ESCRT-dependent and independent pathways, we employed HGS knockdown using two distinct DsiRNA as well as the MA and GW4869 inhibitors targeting exosome biogenesis. The efficiency of HGS knockdown in HeLa cells was monitored by quantitative real-time polymerase chain reaction (qRT-PCR) 72 hours post DsiRNA transfection (Figure 3b). As depicted in Figures 3c and 3d, the depletion of HGS and the administration of MA and GW4869 resulted in a significant reduction in glyco-modified exosomal RNAs, evidenced by a decrease in the IR-800 signal. To reinforce the assertion that this decrease in signal stemmed from the inhibition of exosomal release, we also examined cellular RNAs from the same set of HGS knockdown experiments from Figure 3c. As anticipated, a corresponding increase in cellular glycoRNAs (Figure 3e) was observed, indicating a block in exosome release and corresponding accumulation of exosome and its glycoRNA cargo within the cytoplasm. In summary, our findings underscore the intricate involvement of both ESCRT-dependent and independent pathways in the biosynthesis of glycoRNAs within exosomes.

### Distinctive Signatures of Glyco-Modified Exosomal RNAs Across Human iPSCs, and Induced Neurons

To elucidate the presence of glyco-modified exosomal RNAs across diverse cell types and explore their distinctive signatures we investigated their fate during induced neuronal transdifferentiation (Extended Data Figure 5a). Total Ac4GalNAz-labeled cellular and exosomal RNAs were extracted from human induced pluripotent stem cells (iPSCs) and their derived induced neuron cells (iNs) were subjected to SPAAC reaction after 48 hours of incubation with 100µM Ac4GalNAz-containing media. Importantly, glycolated exosomal RNAs from iPSCs and iNs exhibit a remarkable distinction in their size profile (Extended Data Figure 5b), which was also evident in the cellular RNAs (Extended Data Figure 5c). iPSC-derived exosomes are characterized by the prevalence of slower migrating RNAs, while iNs showcase faster migrating RNAs (sRNAs), aligning with the profiles observed in the more differentiated HeLa and HEK293T cell lines suggesting differentiation dependent distribution of exosomal glycoRNAs.

### Intercellular Communication of Glyco-Modified Exosomal RNAs

Exosome-mediated transfer of RNAs has been shown as a novel mechanism of genetic exchange between cells^27^. To assess the capacity of cells to communicate through the exosome-directed transfer of glycoRNA, naïve HeLa cells were incubated with and without Ac4GalNAz-labeled Hela cell exosomes for 10 and 20 hours (Figure 4a). Following extensive washing to eliminate extracellular exosomes, cellular RNA was isolated and visualized through SPAAC-mediated biotin conjugation. Remarkably, glyco-modified exosomal RNAs were detected in the naïve HeLa cells (Figure 4b and 4c), demonstrating exosome-mediated cellular communication of glycoRNAs.

**Figure 4.**
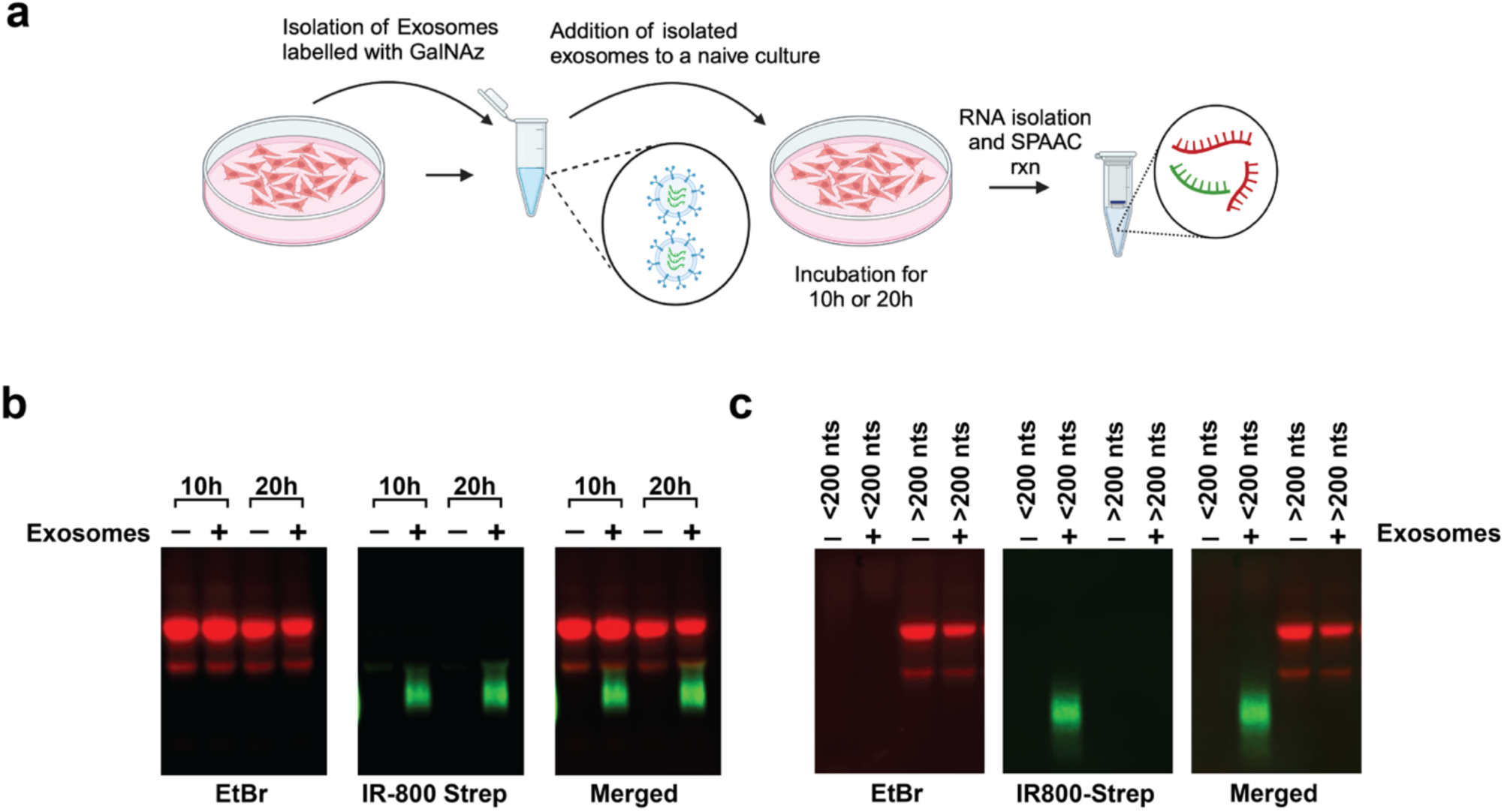
Exosome directed intercellular transfer of glycoRNAs. **(a)** Experimental design overview demonstrating intercellular transfer of glycoRNAs from host cells to naïve cells. **(b)** Total RNA from naïve cells, with and without exposure to Ac_4_GalNaz-labeled exosomes isolated from HeLa cells, was incubated for 10 and 20 hours. After extensive washing to remove extracellular exosomes, total RNA was isolated from the naive cells and subjected to SPAAC-mediated conjugation to visualization the resulting biotin-conjugated glycoRNAs. **(c)** RNA derived from naïve cells subjected to incubation with and without Ac4GalNaz-labeled exosomes isolated from HeLa cells are shown following fractionation into small and large RNA fractions. GlycoRNAs were visualized as in Figure 1.

### Inhibition of Protein Glycosylation curtails Exosomal glycoRNAs

A recent report^28^ has unveiled the amino-carboxypropyl-modified Uridine (acp^3^U) as a crucial site for attaching N-glycans to RNA. Building on this finding and supported by genetic knockout data targeting the DTWD2 enzyme responsible for cellular acp^3^U in mammalian cells, we determined whether acp3U was a primary RNA modification essential for N-glycosylation of RNA in our system. DsiRNA-mediated knockdowns of DTWD2 and another relevant amino-carboxy-propyl transferase, TSR3 were carried out in HeLa cell lines to assess the impact of acp^3^U on Ac4GalNAz-derived glyco-modification. The efficiency of knockdown was evaluated through qRT-PCR (Extended Data Figure 6a). Interestingly, knockdown of either DTWD2 or Tsr3 did not manifest a discernible impact on the levels of Ac4GalNaz-derived glyco-modification (Extended Data Figure 6b and 6c) suggesting the observed RNA glycosylation is not mediated through acp^3^U conjugation.

We next set out to identify the specific sugar precursor responsible for the glyco modification and targeted Galactose-4-Epimerase (GALE) and UDP-Mannose-4-Epimerase (GNE)^29^. GALE and GNE fulfill crucial roles in converting UDP-Glucose to UDP-GalNAc, a process integral to the production of CMP-sialic acid utilized by the Golgi for sialylated glycoconjugation in proteins (Figure 5a). The GalNAc salvage pathway involves GALE as a central enzyme mediating the interconversion between UDP-GlcNAc and UDP-GalNAc. GALE’s enzymatic activity generates key nucleotide sugars essential for glycosylation in the endoplasmic reticulum (ER) and Golgi^29^. Simultaneously, GNE converts UDP-GalNAc and GlcNAc to ManNAc-6 phosphate, a crucial step in glycosylation events shaping cellular processes.

**Figure 5.**
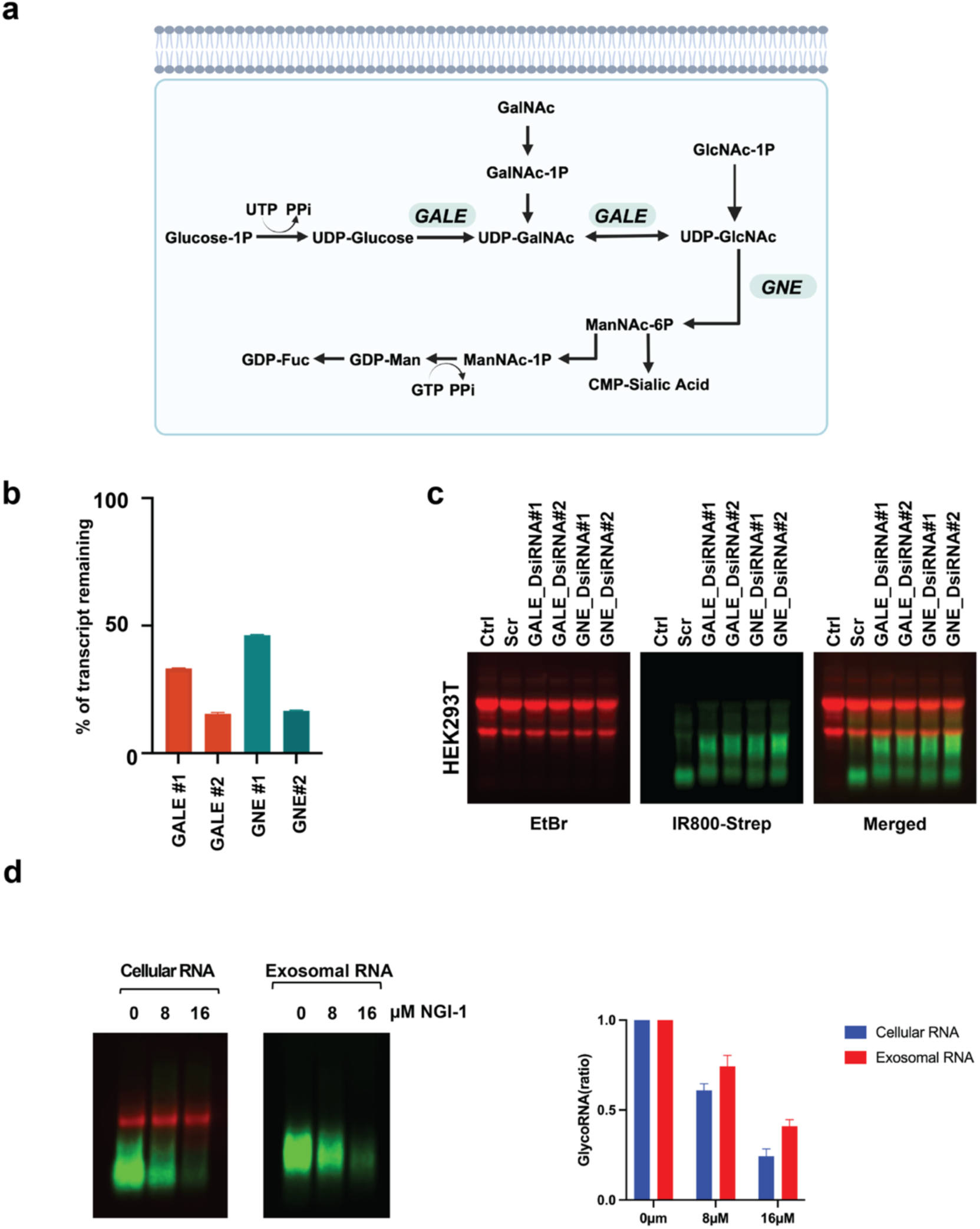
Influence of Steady-State GalNAz Levels on GlycoRNAs. **(a)** The N-acetyl galactosamine (GalNAc) salvage pathway generates UDP-GalNAc, with UDP-galactose 4’-epimerase (GALE) serving as the central enzyme mediating the interconversion between UDP-GlcNAc and UDP-GalNAc. GNE is a pivotal enzyme interconverting UDP-GalNAc and GlcNAc to ManNAc-6 phosphate, a step crucial for glycosylation events shaping cellular processes. **(b)** GALE and GNE depletion by DsiRNA. HEK293T cells were transfected with two distinct DsiRNAs targeting GALE and GNE, followed by a 72-hour incubation. Total RNA extracted from these cells was subjected to qRT-PCR to assess gene knockdown efficiency. **(c)** Analysis of total RNAs from GALE and GNE depleted HEK293T cells. Post-24-hour DsiRNA treatment, 100µM of Ac4GalNaz was introduced, and RNA isolation occurred after 48 hours of incubation with media containing 100µM of Ac4GlcNAz. Following SPAAC-mediated conjugation, the resultant biotin-conjugated glycoRNA was separated on an agarose gel and transferred onto a Nitrocellulose membrane. Visualization was as in Figure 1. **(d)** Inhibition of Oligosaccharyltransferase (OST) by NGI-1. The effect of NGI-1, a specific and potent small molecule inhibitor of OST on glyco-modified RNA production was tested in cells and exosomes. NGI-1 treatment resulted in a dose-dependent loss of Ac_4_GalNaz labelled glycoRNA.

HEK293T cells were transfected with two distinct DsiRNAs targeting GALE and GNE, followed by a 72-hour incubation. Total RNA extracted from these cells was subjected to qRT-PCR to assess the efficiency of gene knockdown (Figure 5b). After 24 hours of DsiRNA treatment, Ac4GalNAz was added to the media and RNA was isolated after 48 hours of incubation, resolved by agarose gel and visualized (Figure 5c). Significantly, the substantial increase in the IR-800 signals following the knockdown of both GALE and GNE strongly suggested an elevated relative steady state of Ac4GalNAz. These observations implicate Ac4GalNAz as the source of the observed modifications. Furthermore, our findings underscore that the steady-state levels of Ac4GalNAz play a pivotal role in determining the abundance of glycoRNAs.

Next, we investigated the impact of NGI-1, a specific and potent small molecule inhibitor of oligosaccharyltransferase (OST)^30^, which transfers branched sugars onto protein in the endoplasmic reticulum, on glycoRNA production. Interestingly, NGI-1 treatment resulted in a dose-dependent loss of glycoRNA labeling for both cellular and exosomal RNAs with Ac4GalNAz (Figure 5d). These data indicate that a block of protein glycosylation also leads to a decrease in RNA glycosylation and a potential coordination between protein and RNA glycosylation. Overall, our results underscore the shared regulatory pathways involving GalNAc and its associated enzymes in orchestrating protein and RNA glyco modifications.

## Discussion

Our investigation into glycoRNAs within exosomes has provided novel insights into the intricate mechanisms of RNA sorting and the potential functional implications of RNA glycosylation. The findings presented in this manuscript align with the emerging understanding of extracellular ss (exRNAs) as multifaceted players in intercellular communication^11,12^. Particularly, the identification of glycoRNAs on the cell surface, as well as the development of membrane-coated nanoparticles for RNA analysis, underscores the complexity and regulatory potential of RNA outside the canonical intracellular environment.

The biogenesis and specific sorting of glycoRNAs into exosomes, as revealed by our study, suggest a targeted mechanism that may be crucial for cell-cell communication and could have implications for the regulation of gene expression. The higher degree of glyco-labeled RNA within exosomes relative to cellular RNAs suggests glycoRNAs are preferentially encapsulated within exosomes. At present, the exact pathway and molecular interactions that facilitate the transport of these glycoRNAs across cellular membranes remain unknown. Nevertheless, considering that glyco-modification of proteins is an important prerequisite for protein sorting into exosomes, it is tempting to speculate that glyco-modification fulfills a similar function for RNA and serves as the conduit for RNA trafficking into exosomes. Moreover, the disruption of oligosaccharyltransferase that transfers branched glycol moieties onto a protein in the ER with the use of NGI-1 inhibitor altering the level of targeted glycoRNAs within exosomes (Figure 5d) indicates accumulation of glycoRNAs to exosomes can be a regulated process. Further detailed understanding of this process could have significant biological and therapeutic potential.

Studies with the NGI-1 inhibitor also reveal a connection between protein and RNA glycosylation. Treatment with NGI-1 to block protein glycosylation resulted in decreased accumulation of exosomal glycoRNAs. This finding indicates that the same pool of glycol moieties is utilized to modify both protein and RNA and implicates a common regulatory pathway. Further studies are necessary to address the potential link between protein and RNA glycosylation and modes of its regulation.

Although we initially set out to identify UDP-GlcNAc capped RNA, two lines of evidence suggest the glycoRNA species detected are likely not glyco-capped RNA but rather consist of internally glyco modified RNA. First, the pleotropic decapping enzyme family of DXO and Rai1 which can decap GlcNAc-capped RNA, do not remove the glyco moiety from the glycoRNA (Figure 1e). Second, the remarkable resistance to RNase A (Figure 1d) and susceptibility to other nucleases (RNase I and micrococcal nuclease) strongly suggests internal modification of the RNA that impedes nucleases either indirectly by affecting the structure or directly by virtue of its modification of the RNA. Its enhanced resistance to nucleases is also indicative of a potential function for glycosylation in the stability of the modified RNA. A detectable resistance to RNAse A treatment was also recently reported for cell surface RNAs detected following Ac4ManNAz treatment of cells ^31^ indicating a common feature of the glyco modification to conferring enhanced nuclease resistant and stability onto an RNA.

The finding in this report complement the recent findings that identified glycoRNA derived from Ac4ManNAz that appear as slow migrating glycoRNAs on an agarose gel and can reside on the cell surface^10,11^. Importantly, the Ac4GalNAz glycoRNA reported here appears to be distinct and considerably more abundant with the assay parameters used (Figure 1b). Moreover, the Ac4ManNAz-derived glycoRNAs are predominantly on the cell surface^28^ while the Ac4GalNAz glycoRNA we detect in the exosome are resistant to nuclease treatment of the intact cell as well as the intact exosome and appear intraluminal. These findings indicate there are at least two classes of glycoRNAs.

In conclusion, our findings contribute to a growing body of evidence that positions extracellular RNAs, and specifically glycoRNAs, as significant entities in the extracellular milieu. The role of Ac4GalNAz-derived RNA glycosylation in the sorting and stabilization of exRNAs within exosomes opens new avenues for research and potential clinical applications. Moreover, the potential of glycoRNAs to serve as biomarkers for disease diagnosis and treatment is particularly tantalizing. Future studies will be essential to unravel the precise mechanisms governing RNA transport across membranes and the functional role of glycoRNAs in exosomes.

## Extended Figures

**Extended Data Figure 1.**
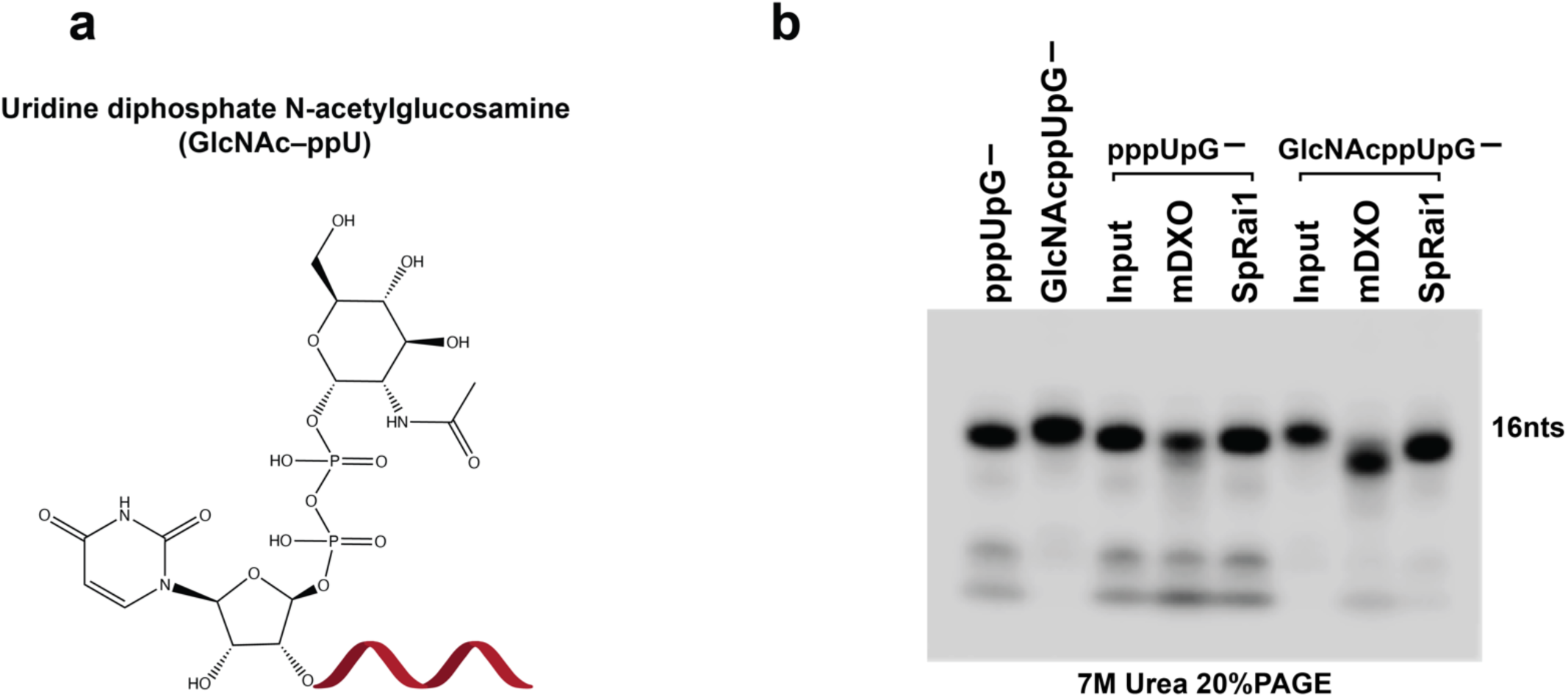
Screening of UDP-GlcNAc capped decapping RNAs. Screening of UDP-GlcNAc capped RNA decapping. **(a)** Schematic representation illustrating the structure of UDP-GlcNAc-capped RNA. **(b)** Analysis of reaction products from *in vitro* decapping assays conducted with 25 nM recombinant mDXO and SpRai utilizing uniformly ^32^P-labeled triphosphate (pppUpG) or UDP-GlcNAc-capped (GlcNAcppUpG) RNA (16 nucleotides) as substrates, denoted accordingly. The resultant products were separated via electrophoresis on 20% polyacrylamide gels containing 7 M urea.

**Extended Data Figure 2.**
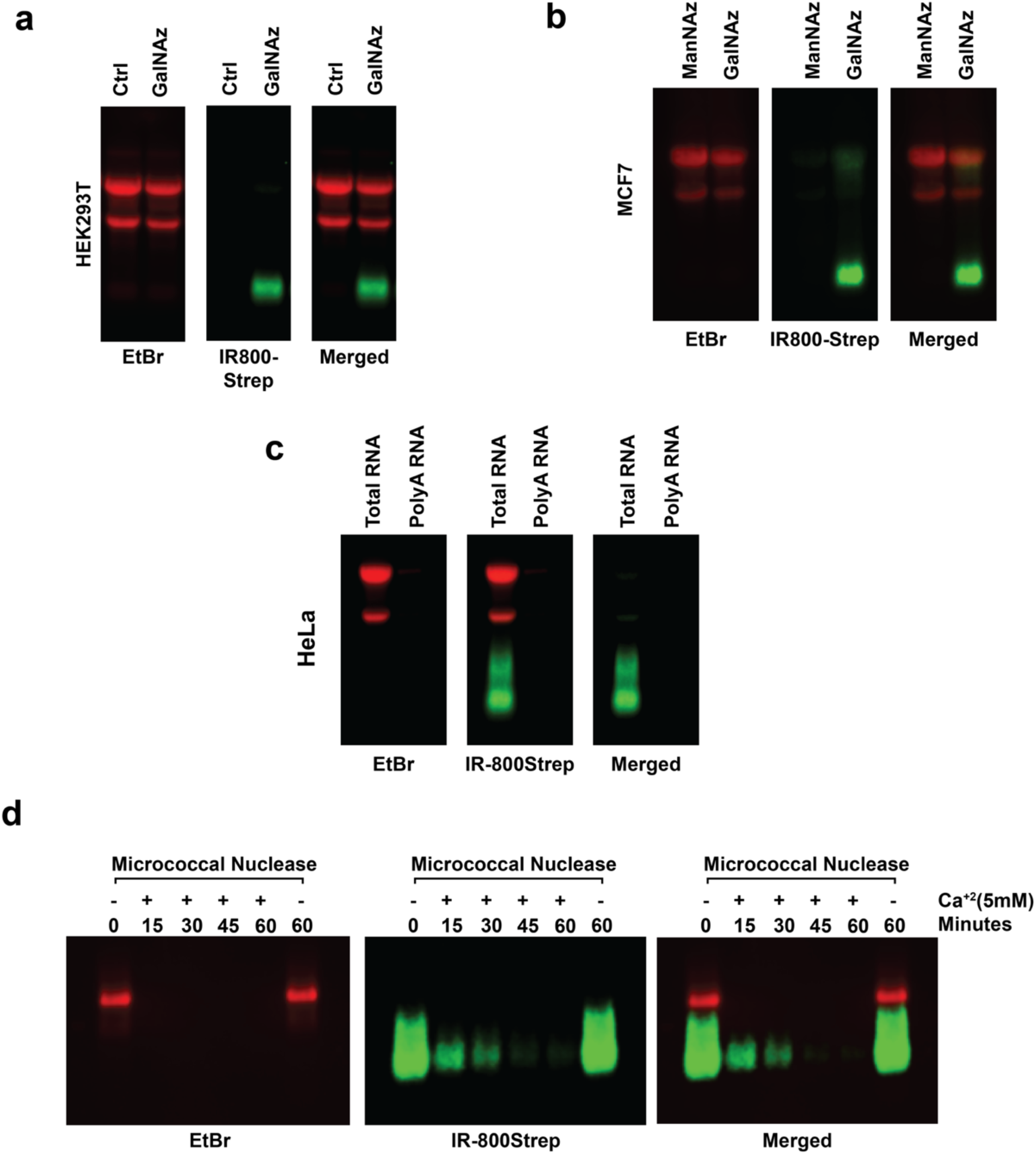
Analysis of Glyco-RNAs in HEK293T and MCF7 cells and impact of RNase treatment. Total RNA from **(a)** HEK293T or **(b)** MCF7 cells obtained following 48 hours of incubation with media containing 100µM of Ac4GalNAz or Ac4ManNAz, underwent SPAAC reaction to attach biotin moieties. The resulting biotinylated RNAs were separated on an agarose gel and transferred onto a Nitrocellulose membrane. Detection was carried out using near-infrared (IR) IRDye® 800CW streptavidin-based labeling for glyco-modified RNAs and ethidium bromide staining for total RNA visualization, employing the Odyssey Fc imaging system (Li-Cor Biosciences) as in Figure 1. **(c)** Following 48 hours of incubation with media containing 100µM of Ac4GalNAz, the enrichment of labeling for both total RNA and Poly A+ RNA was performed using the Poly(A)Purist™ MAG Kit (Invitrogen). The resulting fractions were assessed as described above. **(d)** Time course analysis at the indicated time of 15U micrococcal nuclease digestion of glycoRNAs at 37°C with or without the addition of the Ca^2+^, an essential cation for micrococcal nuclease activity.

**Extended Data Figure 3.**
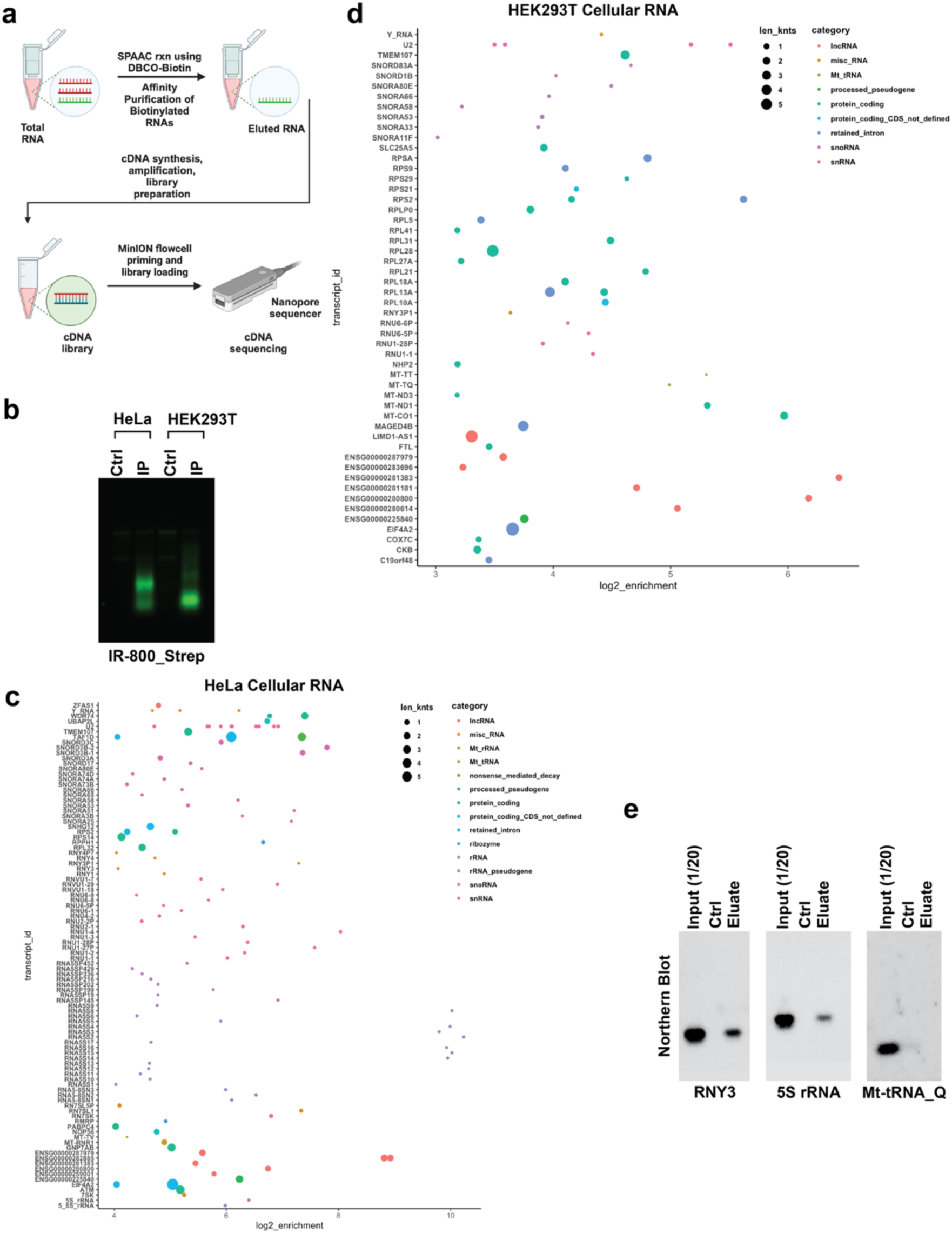
Affinity purification of glycoRNA and Oxford Nanopore sequencing. **(a)** Schematic representation illustrating the process of labeling and affinity purification of glyco-modified RNAs. **(b)** SPAAC conjugated biotinylated small RNA (15 µg) were affinity-purified biotinylated RNA, eluted from streptavidin beads and visualized as in Figure 1. **(c)** Scatter plot analysis depicting Ac4GalNAz-derived labeled RNAs from HEK293T cells and **(d)** HeLa cells. The graphs are color-coded based on RNA categories, and each dot’s size corresponds to the respective RNA’s length. **(e)** Northern blot analysis of RNA eluates obtained from affinity-purified glycoRNAs that were subjected to SPAAC from HeLa cells.

**Extended Data Figure 4.**
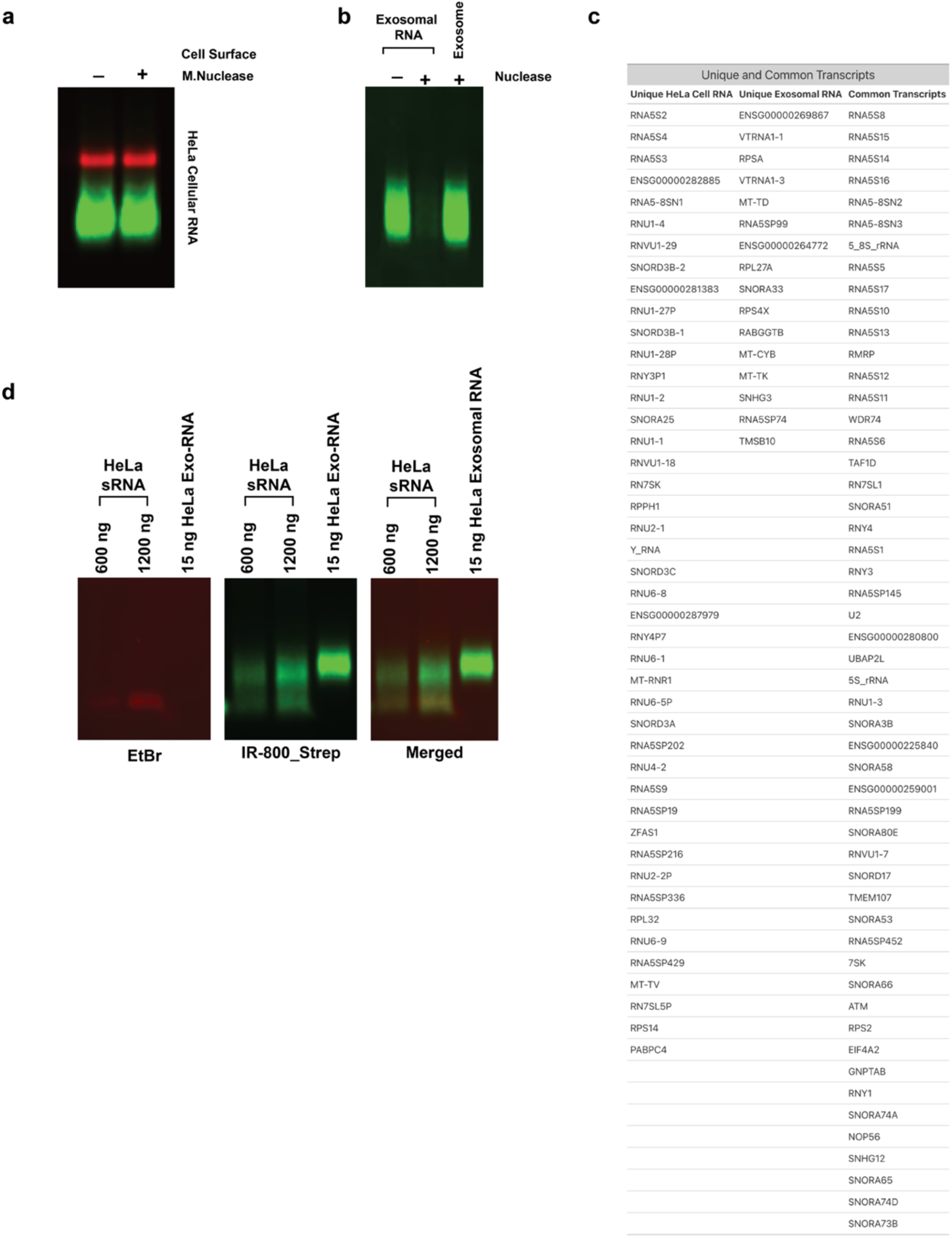
Exosomal glycoRNA. **(a)** Intact HeLa cells or **(b)** intact exosomes isolated from Hela cell cultures, were incubated for 48-hour with 100µM Ac4GalNaz and subjected to micrococcal nuclease. GlycoRNAs subsequently isolated from the treated cells or exosomes were detected. **(c)** List of unique and common transcripts in the cellular and exosomal RNA of HeLa cells enriched over 8-fold (see Figure 2f). **(d)** SPAAC reaction of the indicated amounts of HeLa small RNAs, and exosomal RNAs. The biotin-conjugated RNAs were visualized as in Figure 1

**Extended Figure 5.**
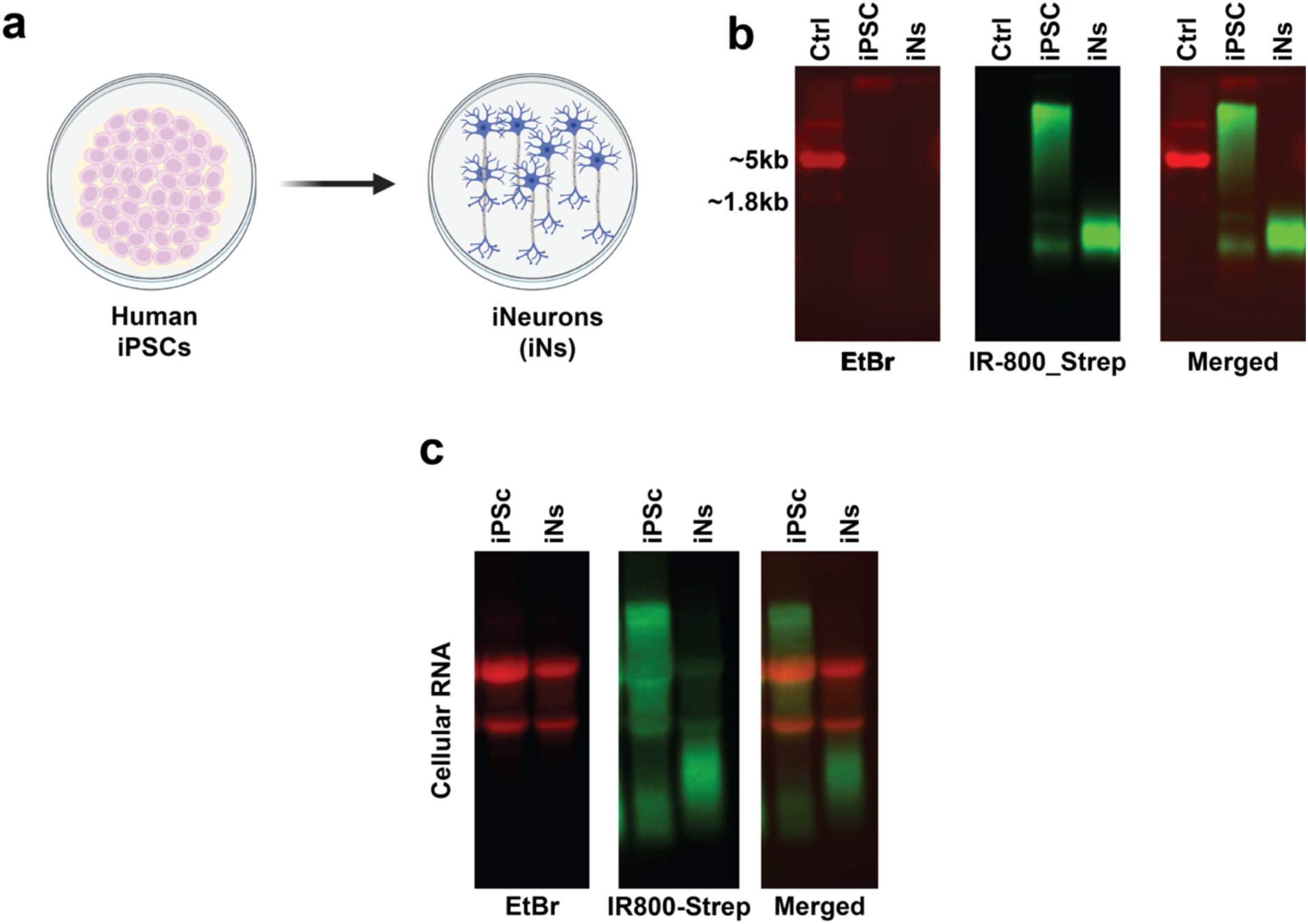
Different patterns of glycoRNAs in iPSC and iN cells. **(a)** Overview of induced neuronal transdifferentiation. **(b)** Exosomal RNAs from human iPSCs and their derived induced neurons (iNs) collected after 48 hours of incubation with 100µM Ac4GalNAz-containing media underwent SPAAC reaction for biotin conjugation. **(c)** Total RNA extracted from human (iPSCs) and their derived induced neurons (iNs) were harvested following a 48-hour incubation with media containing 100µM Ac4GalNaz. Biotin-conjugated glycoRNAs were visualized as in Figure 1.

**Extended Data Figure 6.**
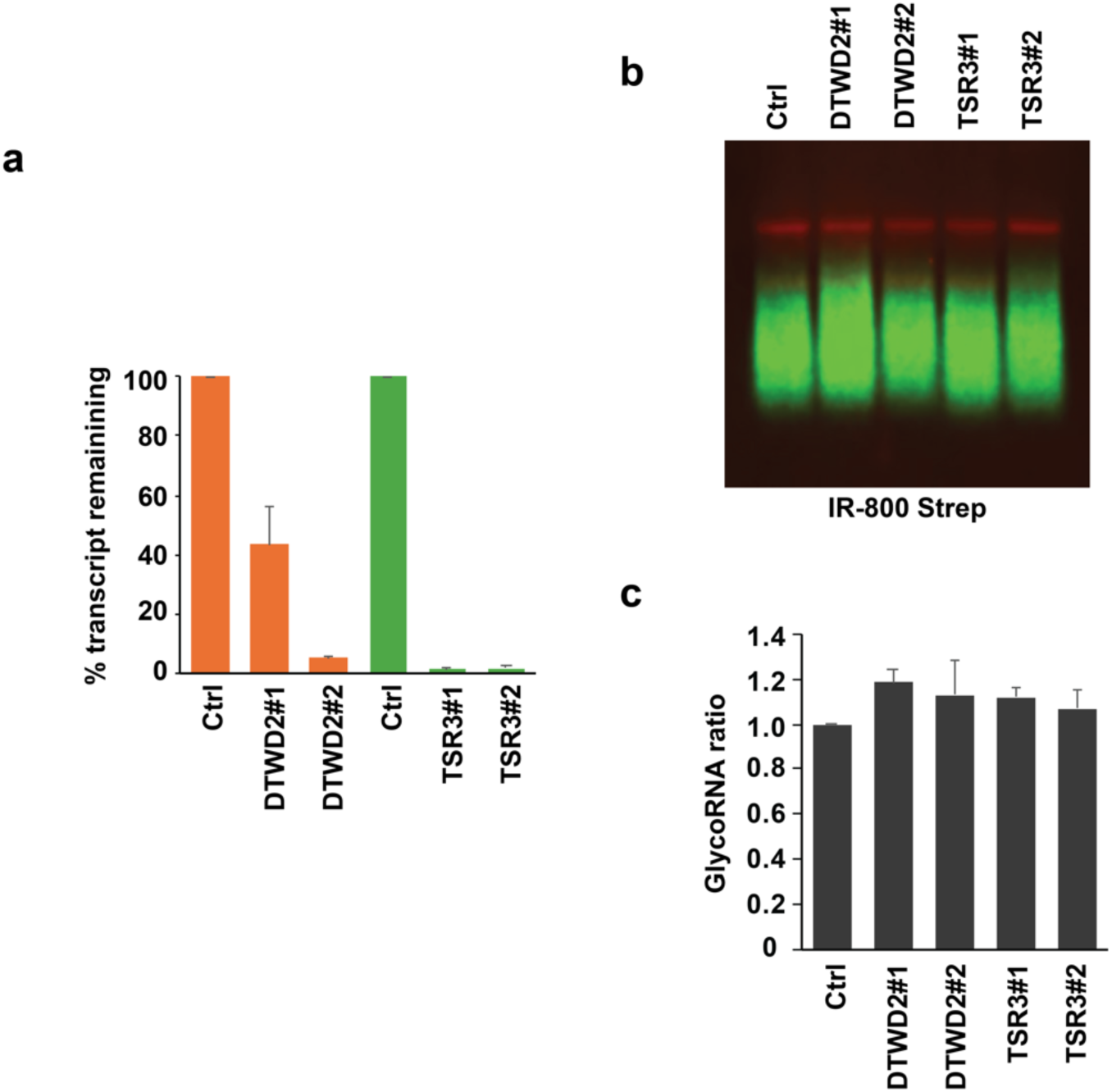
Glycan addition is not mediated through amino-carboxypropyl-modified Uridine (acp^3^U). DTWD2 and TSR3 were knocked down in Hela cells with specific DsiRNAs. After 24 hours DsiRNA treatment, 100µM Ac4GalNAz was added to the medium. Cells were harvested 48 hours after the addition of 100µM Ac4GalNAz. **(a)** The efficiency of target gene knockdown was tested utilizing qRT-PCR analysis. **(b)** Azide-labeled glycoRNA from the control and corresponding knockdowns were subjected to biotin-conjugated SPAAC analysis and visualized as in Figure 1. **(c)** Quantitation of the glycoRNAs in (b) using Image J. GlycoRNA levels in control knockdown were set as 1. Error bars show the +/-SD from three independent experiments (*n* = 3 replicates).

## Methods

### Synthesis of 3’-(guanyl-5′-yl)-uridine 5′-diphosphate N-acetylglucosamine, NAcGlcppUpG, capped RNAs via In Vitro Transcription

To generate RNAs capped with UDP-GlcNAc, we employed *in vitro* transcription using synthetic double-stranded DNA templates T7-GlcNAc-25nt and T7-GlcNAc-40nt. Due to the challenge of incorporating uridine-containing caps, we utilized a 5’-sugar-modified uridine-guanosine (UpG) dinucleotide^32^ – NAcGlcppUpG. This strategy was necessitated by the strong preference of T7 RNAP for transcription initiation with guanosine, which interacts with the N-7 and O-6 of the purine ring^33^. While adenosine-containing cofactors can partially mimic these interactions and initiate transcription^34^, pyrimidine nucleotides like UDP-GlcNAc do not facilitate efficient transcription initiation. Following *in vitro* transcription at 37°C overnight using the HiScribeTM T7 High Yield RNA Synthesis Kit (New England Biolabs (NEB)), RNA purification was performed employing Monarch®RNA Cleanup Columns (NEB) according to the manufacturer’s protocol. For ^32^P-labeled RNAs, the transcription reactions were performed in the presence of [α-^32^P] ATP.

### *In vitro* Decapping Assays

Recombinant Nudix proteins, SpRai1 and mDXO with an N-terminal 6 × His-tag, were purified using His Bind Resin (Novagen). ^32^P-labeled NAcGlcppUpG-cap or pppUpG capped RNAs were incubated with 50 nM of recombinant Nudix proteins and 25nM of SpRai1 and mDXO in decapping buffer (10 mM Tris–HCl [pH 7.5], 100 mM KCl, 2 mM DTT, 2 mM MgCl_2_, and 2 mM MnCl_2_) at 37°C for 30 min. Reactions were terminated by heating at 90°C for 2 min. Decapped products were separated via electrophoresis on 7M urea 20% polyacrylamide gels, visualized using an Amersham Typhoon Biomolecular Imager (GE Healthcare).

### Cell Culture

All cell lines were cultured at 37°C with 5% CO2. Specific cell types used in this study included HeLa, HEK293T, MCF7, and human-induced human pluripotent stem cells (iPSCs)^35^. HeLa, HEK293T, MCF7 cell lines were maintained in DMEM media supplemented with 10% exosome free-fetal bovine serum (FBS) and 1% penicillin/streptomycin (P/S). iPSCs were maintained in mTeSR™ hPSC medium (STEMCELL Technologies).

### Metabolic Labeling with Ac4GalNAz

Metabolic labeling of cells was achieved using N-azidoacetylgalactosamine-tetraacylated (Ac4GalNAz) or N-azidoacetylmannosamine-tetraacylated (Ac4ManNAz) or N-azidoacetylglucosamine-tetraacylated (Ac4GlcNAz). Cells were cultured under standard conditions and treated with 100 µM Ac4GalNAz for specified durations. Following treatment, cells were harvested for subsequent RNA analysis.

### RNA Isolation and treatment

Total RNA was extracted from treated and control cells using TRIzol reagent according to the manufacturer’s instructions. The RNA was then treated with DNase I to remove any potential DNA contamination. To ensure the removal of proteins, samples were subjected to Proteinase K treatment and purified again using the Monarch®RNA Cleanup Columns (NEB) according to the manufacturer’s protocol. To enrich for polyadenylated RNA, total RNA purified as above was used as the input for the Poly(A) Purist MAG Kit (Thermo Fisher Scientific) following the manufacturer’s protocol.

### Strain-promoted alkyne azide cycloaddition (SPAAC)

To conjugate biotin to Ac4GalNAz-labeled sugars, a copper-free click chemistry approach was employed using dibenzocyclooctyne (DBCO)-PEG4-biotin (MilliporeSigma). The SPAAC reactions were performed following the previously established protocol^18^. Here 25µg of Ac4GalNAz-labeled RNA-bearing azide RNA in pure water were mixed with 1x volumes of MK-Gel Loading Buffer (90% Formamide, 15mM EDTA, and 0.025% SDS) and 500 μM DBCO-biotin (MilliporeSigma). Typically, these reactions consisted of 20 μL MK-Gel Loading Buffer, 18 μL RNA (from ADPRC reaction), and 2 μL of a 10mM DBCO-biotin reagent stock solution. The reactions were carried out at 55°C for 15 minutes to process the copper-free glycoRNA Click, halted by adding 310 μL water, followed by purification of the conjugated RNA using Monarch®RNA Cleanup Columns (NEB). The RNA underwent analysis either through gel electrophoresis or affinity purification as detailed below. To confirm that RNA molecules were labeled following the SPAAC reaction, the RNA was subjected to RNAse A (1µg), RNase I (10U), micrococcal nuclease (15U) or DNase I (2U) at 37°C for 1 hour before loading onto the gel as described below.

### RNA gel electrophoresis, capillary blotting, and near-infrared fluorescent imaging

For blotting analysis of either enriched RNA following biotin-mediated affinity purification or directly after the SPAAC reaction, RNA was heat-incubated at 90°C for 2 minutes in formaldehyde loading buffer (FLB) (50% Formamide, 6% formaldehyde, 50mM HEPES, pH 7.8, 0.5µg/mL ethidium bromide, and 10% glycerol). Samples were loaded onto a 1.2% agarose gel and electrophoresed at 120V for 1 hour. Before transferring the gel onto a 0.45 μm nitrocellulose membrane (Cytiva), total RNA was visualized using a UV gel transilluminator. RNA transfer occurred overnight via passive; upward transfer facilitated by the capillary flow of the 10X SSC buffer. Following the transfer, RNA was cross-linked to the NC using UV-C light (0.2 J/cm2). Subsequently, the membrane was blocked with Odyssey Blocking Buffer (Li-Cor Biosciences) for 30 minutes at room temperature (RT). After blocking, IRDye® 800CW streptavidin (Li-Cor Biosciences) was diluted to 1:7,000 in Odyssey Blocking Buffer and used to stain the NC membrane for 30 minutes at RT followed by three washes with PBST (ResearchProductInternational). Before scanning, the membranes were briefly rinsed in 1x PBS and scanned by an Odyssey Fc (Li-Cor Biosciences) with the software set to auto-detect the signal intensity for both the 600 and 800 channels. The 600 channel was utilized for ethidium bromide, while the 800 channel was utilized for IRDye® 800CW streptavidin detection.

### Subcellular Fractionation

To obtain highly purified nuclei and nuclei free cytoplasm, we employed a commercially available NE-PER® (Thermo Scientific) Nuclear and Cytoplasmic Extraction Kit for efficient cell lysis and extraction of separate cytoplasmic and nuclear protein fractions following the manufacturer’s instruction. Briefly, Ac4GalNAz labeled HeLa cells were harvested using trypsin-EDTA washed with PBS, and pelleted. After discarding the supernatant, the cell pellet was treated with ice-cold cytoplasmic extraction reagent I (CER I). Cytoplasmic and nuclear proteins were extracted by vortexing the cell suspension, adding CER II, and centrifuging to collect the cytoplasmic fraction. The nuclear pellet was then suspended in nuclear extraction reagent (NER), vortexed, and centrifuged to obtain the nuclear extract. One third of this fraction was used for the Western Blotting whereas the rest was used to extract RNA using a TRIzol reagent.

For cytosolic and crude membrane fraction isolation from the Ac4GalNAz -labeled HeLa cells, we utilized a commercial membrane protein extraction kit - The ProteoExtract® Native Membrane Protein Extraction Kit (EMD Millipore). This extraction method involves sequential lysis steps, initially releasing soluble cytosolic proteins and RNAs, followed by the disruption of membranous organelles such as the plasma membrane, Golgi, and ER. In brief, cultured HeLa cells underwent media removal, followed by two washes with ice-cold Wash Buffer. Subsequently, Extraction Buffer I (containing protease inhibitor) was applied to the cells and incubated for 10 minutes at 4°C with gentle rocking, yielding the ‘cytoplasm’ fraction. This was followed by the addition of Extraction Buffer II (also supplemented with protease inhibitor) to the cells, which were further incubated for 30 minutes at 4°C with gentle rocking, resulting in the ‘ER/membrane’ fraction. Here again, 1/3^rd^ of the fraction was used for Western Blotting whereas 2/3^rd^ of it was used for s extraction using TRIzol extraction method.

### Nanopore-Based RNA Sequencing

Five micrograms of small RNA were isolated from biological replicates of both HE29T, HeLa cells and HeLa exosomes and polyadenylated in a 25 μL reaction mix containing 2 μL of 10X E. coli poly(A) polymerase buffer, 2 μL of 10 mM ATP, 15-17 μL of nuclease-free water, and 1 μL of E. coli poly(A) polymerase (5 U/μL) (NEB), with incubation at 37°C for 2 minutes. To stop the reaction, 5 μL of 50 mM EDTA was added to reach a final concentration of 10 mM EDTA. Following incubation, 45 μL of RNase-free SPRI beads were added to the reaction and mixed on a mixer for 5 minutes at room temperature. The sample was then centrifuged, and the supernatant discarded after magnetic separation. The beads were washed twice with 200 μL of freshly prepared 70% ethanol, and any residual ethanol was removed by air-drying the beads for 30 seconds. The beads were resuspended in 15 μL of nuclease-free water and incubated on ice for 5 minutes. The sample was cleared by magnetic separation, and the 15 μL eluate containing the polyadenylated RNA was retained.

Following the SPAAC reaction, enrichment was achieved through specific binding to streptavidin-coated magnetic beads, as detailed previously^18^. Briefly, RNA samples were incubated at 25°C for 30 minutes with 15 μL of pre-blocked magnetic Dynabeads™ MyOne™ Streptavidin T1. Pre-blocking was carried out with 100 ng/µL of bacterial small RNAs in 100 μL of immobilization buffer, comprising 10 mM Tris-HCl (pH 7.5), 1 mM EDTA, and 2 M NaCl. Following thorough washing with a wash buffer (10 mM Urea, 5 mM Tris-HCl [pH 7.5], 0.5 mM EDTA, and 1 M NaCl) for five cycles at 25°C for 5 minutes each, biotinylated RNAs were eluted by incubating the beads with 20 μL of MK-Gel Loading Buffer at 90°C for 2 minutes. Subsequently, the biotinylated RNAs underwent purification using Monarch® RNA Cleanup Columns (NEB) as outlined above and were eluted in 10 µL of nuclease-free water.

Nanopore RNA sequencing and subsequent bioinformatics analysis were carried out using a comprehensive pipeline. Nanopore sequencing libraries were prepared according to the manufacturer’s instructions using the PCR-cDNA Barcoding Kit (SQK-PCB111.24) with R9.4.1 flow cells. Raw nanopore reads underwent quality control using NanoQC (https://github.com/wdecoster/nanoQC) to ensure data reliability. Subsequently, alignment to the reference genome (https://www.gencodegenes.org/human/release_43.html) was performed using Minimap2 (https://github.com/lh3/minimap2#uguide), resulting in the generation of a Binary Alignment Map (BAM) file.

Transcript quantification was performed using Salmon (version 1.10.1) with the aligned reads as input. The resulting quantification files (e.g., transcripts per million, TPM) were generated for downstream analysis. For differential expression analysis, edgeR (version 3.36.0) was employed. The quantification files from Salmon were converted to count matrices, which served as input for edgeR. The count data were normalized using the trimmed mean of M-values (TMM) method. Visualization of transcriptomic data was conducted using the ggplot2 package in R.

### Exosome Isolation

Extracellular vesicles, including exosomes, were isolated using the Total Exosome Isolation Reagent (Invitrogen^TM^). A total of 50 mL of cell culture media harvested from HeLa cell cultures underwent exosome isolation following a standardized protocol. Initially, the harvested media underwent triple centrifugation: first at 300 × g for 10 minutes, then at 2000 × g for another 10 minutes, and finally at 10,000 × g for 30 minutes at 4°C, aimed at effectively removing cells and debris. The resulting supernatant was carefully transferred to new tubes. Exosomes were isolated from the cell-free culture media using the Total Exosome Isolation (from cell culture media) reagent according to the manufacturer’s instructions. Briefly, 50 mL of cell-free culture media was mixed with 0.5 volumes of the isolation reagent, vortexed thoroughly, and incubated overnight at 4°C. After incubation, samples were centrifuged at 10,000 × g for 1 hour at 2°C, and the supernatant was aspirated and discarded, leaving the exosome-containing pellet. Pellets were then resuspended in 400 µL of 1X PBS. The isolated exosomes were subsequently subjected to RNA isolation using the TriZol extraction method and Western blot analysis using the exosome marker CD9 to confirm the presence of exosomal markers.

### Intercellular Transfer of GlycoRNAs via Exosomes

To investigate the transfer of glycoRNAs via exosomes, Ac4GalNaz-labeled exosomes were isolated from HeLa cells cultured in two 150 mm plates containing 30 mL of DMEM medium, as described above. The isolated exosomes were then incubated with naïve cells for 10 and 20 hours. Following incubation, the cells underwent extensive washing to remove any extracellular exosomes. Total RNA was subsequently extracted from the naïve cells and subjected to strain-promoted azide-alkyne cycloaddition (SPAAC)-mediated conjugation to enable visualization of biotin-conjugated RNAs.

### DsiRNA-mediated knockdown and NGI-1 treatment

HeLa cells were seeded 18 hours prior to transfection to achieve an optimal ∼50% confluency in a 10cm plate. Before transfection, each well was supplemented with 1.0 mL of complete medium containing serum and antibiotics, followed by an incubation period of 30 to 60 minutes. DsiRNA, at a final concentration of 40nM per plate, was diluted into 800 µL of PepMute™ Transfection Buffer working solution and thoroughly mixed. Subsequently, 20 µL of PepMute™ Plus reagent was added and thoroughly mixed by pipetting. The mixture was then incubated at room temperature for approximately 15 minutes to facilitate the formation of the transfection complex. The transfection mix was carefully added drop-wise to the cells, with gentle rocking of the plate, before being returned to the CO2 incubator. After 24 hours, 100µM of Ac4GalNAz was added to each plate and the cells were incubated for another 48 hours. Gene silencing was assessed 72 hours post-transfection, and the labeled RNA was assessed by blotting as described above. In the case of NGI-1, Manumycin, or GW4869 treatment, cells were treated with the respective compounds for 24 hours. Following this treatment, Ac4GalNAz was added, and the cells were incubated for an additional 48 hours before harvesting either the cells or the media to assess their impact on intracellular or extracellular RNA labeling, respectively.

### Inducing Neuronal Transdifferentiation from Induced Pluripotent Stem Cells

Human iPSCs were cultured on Matrigel-coated plates in mTeSR™ medium (STEMCELL Technologies) and directly differentiated into induced neurons (iNs) using an inducible lentiviral system as previously described^35^. For neuronal transdifferentiation, iPSCs were dissociated into single cells using Accutase and seeded onto Matrigel-coated supplemented with Y-compound (5 µM) and transduced with lentiviruses encoding rtTA (FUW-M2rtTA) and NGN2 (Tet-O-Ngn2-puro). Transfected cells were grown in neurobasal culture medium containing B27 supplement (Invitrogen). Doxycycline (2 µg/ml) was added 24 hrs post-transfection to induce NGN2 and infected cells were selected with the addition of puromycin (2 µg/ml) for 48 hrs. Cells were cultured for 7 days in neurobasal culture medium supplemented with B27 and doxycycline. During the transdifferentiation process, cells were treated with 100 µM Ac4GalNAz for 48 hours on day 7 of induction. After this labeling period, the cells were washed with PBS and harvested for RNA extraction. Total RNA was extracted using TriZol. The labeled RNA was subjected to copper-free biotin-azide click chemistry to attach a biotin moiety to the labeled RNA molecules and the biotinylated RNA was visualized using IRDye® 800CW streptavidin detection.

## Data availability

All unique materials and reagents generated in this study are available from the corresponding author with a completed material transfer agreement. The sequencing data is deposited at NCBI Sequence Read Archive (SRA) submission: SUB14589150. (Reviewer’s link: https://dataview.ncbi.nlm.nih.gov/object/PRJNA1132718?reviewer=viqljgj42hcgakgl9p3j511jhc). Any additional information required to reanalyze the data reported in this paper is available from the corresponding author upon request. All data are available in the main text or supplemental material.

### Acknowledgement

We are grateful to Dr. Joanna Kowalska (University of Warsaw) for generously providing the NAcGlcppUpG glyco-dinucleotide. This work was supported by National Institutes of Health (NIH) grant GM149262 (M.K.).

## Author Contributions

M.K., S.S. and X.J. designed the experiments. J.Y. carried out the iPSC experiments in Extended Data Figure 5. X.J. carried out the experiments in Figures 1b, 1d, 5d, Extended Figures 2d, 4a and 6. All other experiments were by S.S.. S.S. and M.K. wrote the manuscript with input from all authors.

